# Immune tolerance of human induced pluripotent stem cell-derived myogenic progenitor cells in humanized mice

**DOI:** 10.1101/550699

**Authors:** Basma Benabdallah, Cynthia Désaulniers-Langevin, Marie-Lyn Goyer, Chloé Colas, Chantale Maltais, Yuanyi Li, Jean V. Guimont, Jacques P Tremblay, Elie Haddad, Christian Beauséjour

## Abstract

It is still unclear if immune responses will compromise the large scale utilization of cell therapies derived from human induced pluripotent stem cells (hiPSCs). To answer this question, we used humanized mouse models generated by the adoptive transfer of peripheral blood mononuclear cells (Hu-AT) or the co-transplantation of hematopoietic stem cells and human thymic tissue (Hu-BLT). Using these mice we evaluated the engraftment in skeletal muscle of myogenic cells either obtained directly from a muscle biopsy or differentiated from hiPSCs or fibroblasts. Our results showed that while allogeneic grafts were rejected and highly infiltrated with human T cells, engraftment of autologous cells was tolerated, indicating reprogramming and differentiation procedures are not immunogenic. We also observed that hiPSC-derived myogenic progenitors are not targeted by autologous T cells and natural killer (NK) cells *in vitro*. Overall, our findings suggest that hiPSC-derived myogenic progenitors will be tolerated in the presence of a competent human immune system.

**SIGNIFICANCE:** The immunogenicity of human iPSC-derived cells will strongly influence their use in regenerative medicine. This important feature has so far been mostly understudied given the necessity to have access to humanized mice reconstituted with an immune system autologous to iPSC-derived cells. Using two distinct mouse models we here provide evidences that human immune cell infiltration in skeletal muscle should not be used as the sole marker to predict immunogenicity. Indeed, we show that human iPSC-derived myogenic progenitors, similar to primary human myoblasts, induced autologous T cell infiltration yet without compromising engraftment. Our study provides essential pre-clinical data supporting the usage of human iPSC-derived myogenic progenitor cells.

## INTRODUCTION

The controversial possibility that hiPSC-derived cells arouse an autologous immune response could compromise their use in clinic. Indeed, mouse iPSCs were shown to be immunogenic when injected in syngeneic recipients ^1, 2^ and lead to the development of mechanisms similar to self-tolerance ^3-7^. Using humanized BLT mice (Hu-BLT), Zhao et al. showed that autologous hiPSC-derived smooth muscle cells are immunogenic while hiPSC-derived retinal pigment epithelial cells were not ^8^. Yet, the extent to which engrafted autologous compared to allogeneic hiPSC-derived cells were tolerated was not evaluated. In addition, the immunogenicity observed towards iPSC-derived smooth muscle cells was based on the infiltration of autologous T cells, and graft function or survival was not assessed. Moreover, since BLT mice are deficient in functional NK cells^9^, the *in vivo* contribution of the innate immunity, particularly the role of NK cells in the immunogenicity of hiPSC-derived cells was not addressed. To this end, using a combination of humanized mouse models (Hu-BLT and Hu-AT), we provide evidences that skeletal muscle engraftment is not impaired despite low level autologous immune cell infiltration.

## METHODS

Detailed methods can be found in the supplementary file.

## RESULTS

### Infiltration of autologous T cells in muscle does not mediate immune rejection in Hu-BLT mice

To evaluate the immunogenicity of skeletal muscle cell grafts, we used myogenic cells either obtained directly from a fetal muscle biopsy (myoblasts) or myogenic progenitor cells (MPCs) differentiated from fetal liver fibroblasts (Figures 1A and 2A). MPCs were generated using a two-step protocol as we previously described ^10^. Myoblasts and MPCs were highly positive for the myogenic markers and fully capable of forming myotubes *in vitro* (Figures S1A and S1B). Immunogenicity was determined after injecting myogenic cells into the *Tibialis anterior* of Hu-BLT mice with either an autologous or allogeneic immune system (Figure S2A-C). Four weeks after transplantation, muscle sections had higher number of human dystrophin-expressing myofibers resulting from the fusion of autologous compared to allogeneic MPCs (Figures 1B and 1C). As expected, muscles transplanted with allogeneic MPCs were highly infiltrated with CD4+ and CD8+ T cells (Figure 1D). Surprisingly, transplanted autologous muscles were also infiltrated with T cells although at a lower level (Figure 1E). Importantly, the engraftment of MPCs was not impaired by the infiltration of autologous T cells as an equal number of fibers expressing the human dystrophin were formed when MPCs were transplanted in non-reconstituted NSG mice (Figure 1C). The injection of the immunosuppressive drug FK506 to Hu-BLT mice prevented the rejection of allogeneic myoblasts, consistent with a T cell-dependent immune rejection mechanism (Figures S2D-F). In support of this observation, allogeneic but not autologous MPCs and myotubes were able to activate T cells *in vitro* after a three day-coculture with splenocytes collected from Hu-BLT mice (Figure 1F).

**Figure 1.**
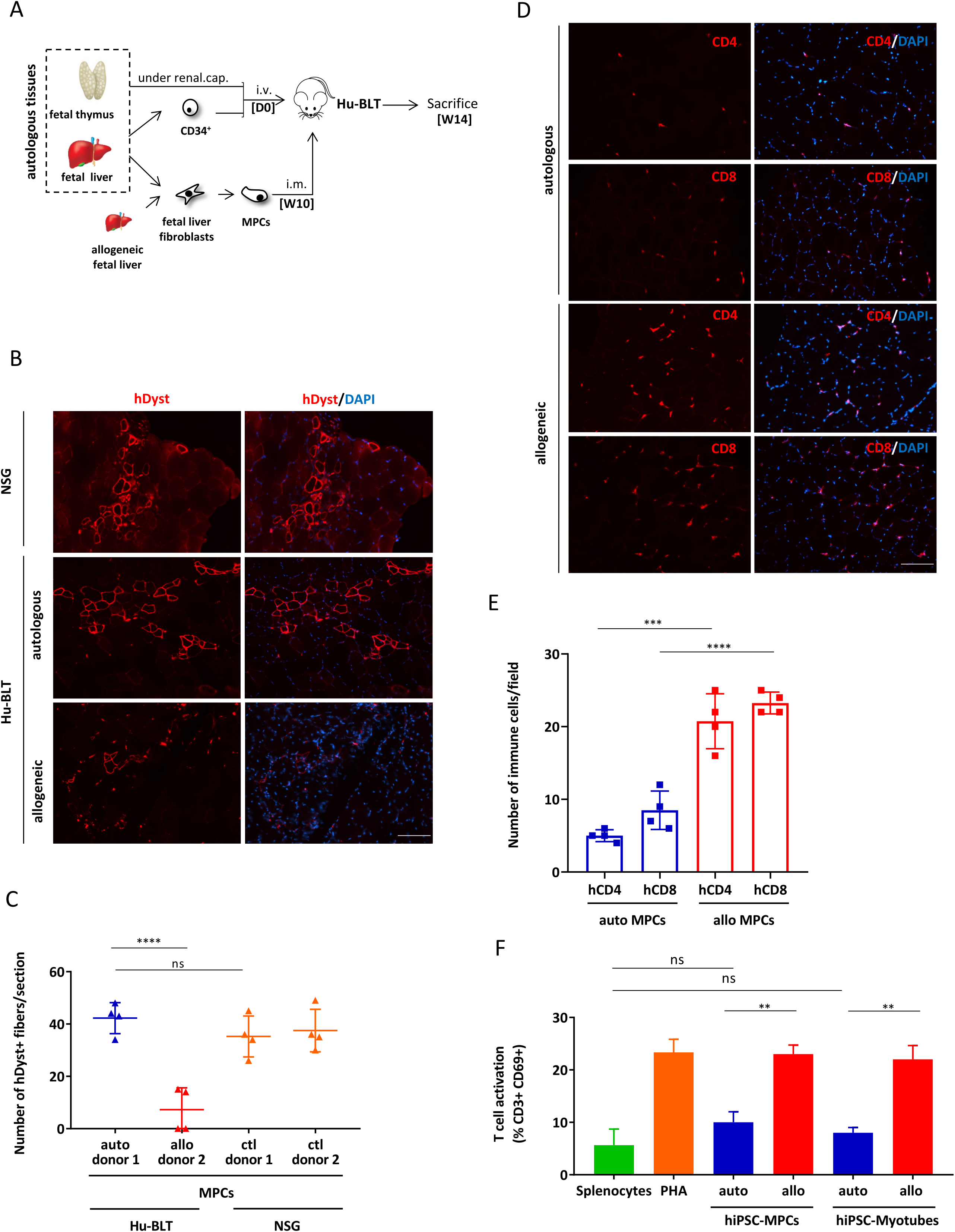
Fibroblast-derived myogenic progenitor cells are immune tolerated in Hu-BLT mice. **(A)** Schematic of the BLT humanized mouse model (Hu-BLT). NSG mice were transplanted with human fetal thymus and autologous liver-derived CD34^+^ cells. At week 10 (W10) post immune reconstitution, human myogenic progenitor cells (MPCs) derived from fetal liver fibroblasts were injected in the skeletal muscle of Hu-BLT. Mice were sacrificed 4 weeks later, and muscles were harvested and analyzed for cell engraftment and infiltration of immune cells. **(B)** Representative photos of muscle sections from NSG or Hu-BLT mice showing increased engraftment of autologous compared to allogeneic human MPC and resulting human specific dystrophin-positive fibers (in red). DAPI staining was performed to visualize nuclei (in blue). Showed are photos taken at 20X. Scale bar, 100 µm. **(C)** Counts of human dystrophin positive fibers observed in muscle sections of Hu-BLT mice transplanted with MPCs as shown in panel **B**. Also showed are counts in non-immune reconstituted NSG mice to assess overall engraftment potential of the different donors. Each dot represents the mean ± SEM number of dystrophin positive fibers from 2-3 randomly selected sections in each muscle (n=4). **(D)** Representative photos showing CD4+ and CD8^+^ T cell infiltration (in red) in muscle sections of Hu-BLT mice transplanted with autologous or allogeneic MPCs. DAPI staining was performed to visualize nuclei (in blue). Showed are photos taken at 20X. Scale bar, 100 µm. **(E)** Frequency of CD4+ and CD8+ T cells infiltrated in muscle sections of Hu-BLT mice as shown in panel D. Each dot represents the mean ± SEM number of T cells in 2-3 randomly selected fields in each muscle (n = 4). **(F)** Splenocytes were collected from Hu-BLT mice 10 weeks after immune reconstitution, and the activation of T cells (as determined by gating CD3^+^ /CD69^+^ cells) was measured after a co-culture with autologous or allogeneic MPCs or myotubes (ratio 1:2) for 3 days. PHA was used as a positive control. Shown is the mean ± SEM of 2 independent experiments done in triplicate using splenocytes collected from 2 different mice.

**Figure 2.**
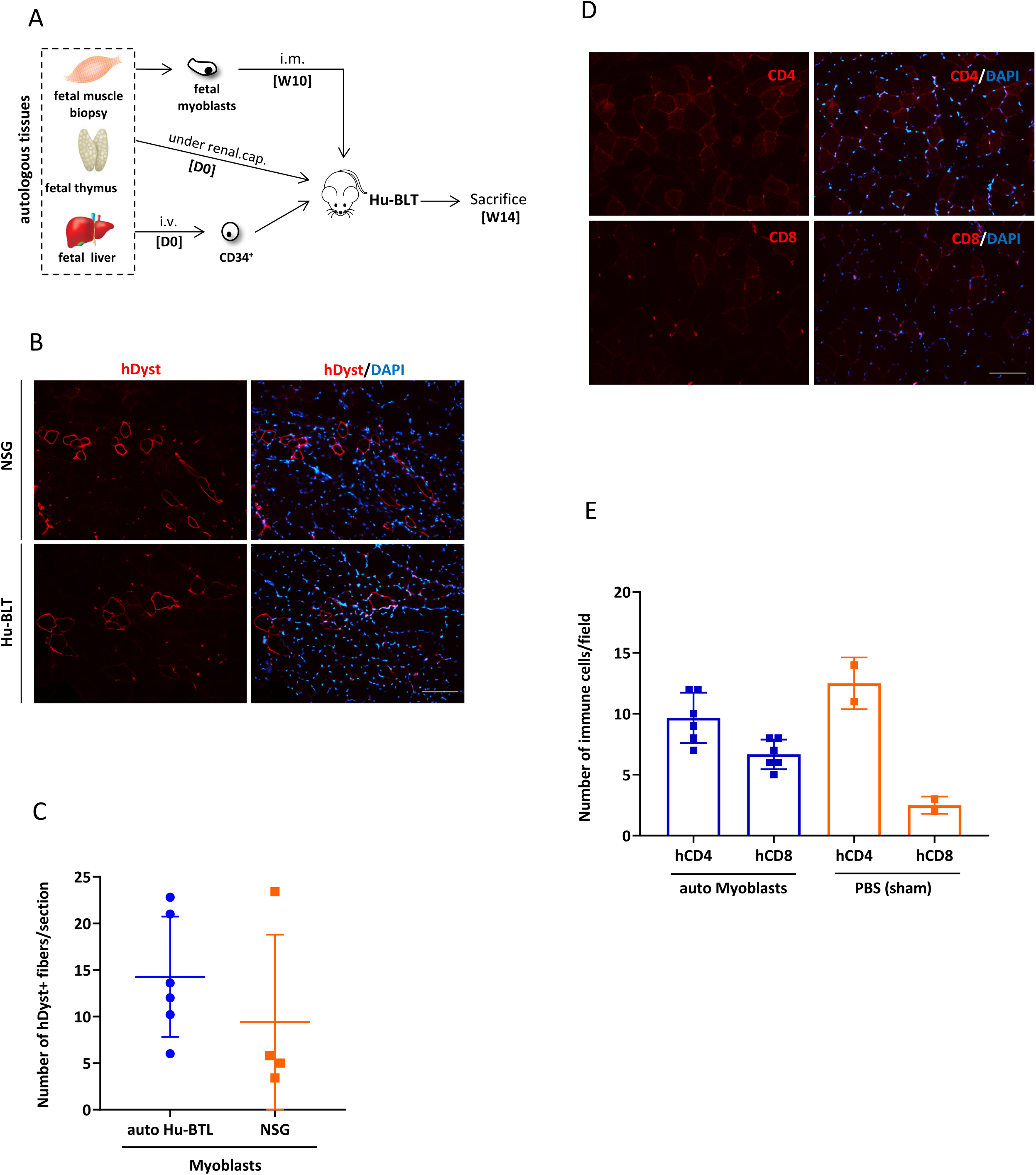
Infiltration of autologous T cells in muscles does not prevent engraftment of fetal myoblasts in Hu-BLT mice. **(A)**Schematic of the BLT humanized mouse model (Hu-BLT). NSG mice were transplanted with human fetal thymus and autologous liver-derived CD34^+^ cells. At week 10 (W10) post immune reconstitution, human myoblasts isolated directly from a fetal muscle biopsy were injected in the skeletal muscle of Hu-BLT. Mice were sacrificed 4 weeks later, and muscles were harvested and analyzed for cell engraftment and infiltration of immune cells. **(B)** Representative photos of muscle sections from Hu-BLT mice showing engraftment of autologous human fetal myoblasts and resulting dystrophin positive fibers (in red). DAPI staining was performed to visualize nuclei (in blue). Showed are photos taken at 20X. Scale bar, 100 µm. **(C)** Counts of human dystrophin-positive fibers observed in muscle sections of Hu-BLT mice transplanted with autologous fetal myoblasts as shown in panel **B**. Also showed are counts in non-immune reconstituted NSG mice to assess overall engraftment potential of the donor. Each dot represents the mean ± SEM number of dystrophin positive fibers from 2-3 randomly selected sections in each muscle (n=4-6). **(D)** Representative photos showing CD4+ and CD8^+^ T cell infiltration (in red) in muscle sections of Hu-BLT mice transplanted with autologous fetal myoblasts. DAPI staining was performed to visualize nuclei (in blue). Showed are photos taken at 20X. Scale bar, 100 µm. **(E)** Frequency of CD4+ and CD8+ T cells infiltrated in muscles of Hu-BLT mice as shown in panel **D**. Each dot represents the mean ± SEM number of T cells in 2-3 randomly selected fields in each muscle (n=2-6).

To determine if the infiltration of T cells was specific to the differentiation procedure of MPCs, we also transplanted muscles of Hu-BLT mice with autologous myoblasts isolated directly from a fetal muscle biopsy (Figure 2A). To our surprise, four weeks after cell transplantation, we found muscle sections were infiltrated with CD4+ and CD8+ T cells at a level similar to what we observed in sham-operated mice or muscles transplanted with autologous MPCs, yet without compromising engraftment (Figures 2B-E). Our hypothesis is that T cells may have been attracted by inflammation-induced damage following the transplantation procedure or by an altered secretory phenotype acquired during the short *in vitro* expansion period of myoblasts. Another possibility is that aberrantly expressed antigens may have attracted and induced tolerance/exhaustion of infiltrating autologous T cells. Unfortunately, the low level of infiltrating T cells in muscle did not allow to test these hypothesis.

### hiPSC-MPCs are not the target of T and NK cells *in vitro*

To better reflect future clinical settings, we then choose to differentiate MPCs from hiPSCs, instead of from fibroblasts, and evaluate their *in vitro* immunogenicity in autologous and allogeneic settings. We first generated hiPSC clones using the integration-free (Sendai virus) approach and confirmed they had normal karyotypes, expressed the classical markers of pluripotency and were able of forming *in vivo* teratoma in immunodeficient mice (Figures S3A, S3B and data not shown). Using the same two-step differentiation protocol we then generated MPCs and showed these cells were positive for MHC-I, costimulatory molecules and NK cell-receptor ligands (Figure S1A and C). We then evaluated the immunogenicity of hiPSC-MPCs in vitro by performing a MLR-like assay using PBMCs and a cytotoxicity assay using PBMC-purified NK cells. Our results show that hiPSC-MPCs and their derived myotubes significantly activated allogeneic but not autologous T cells (Figures 3A and 3B). Conversely, NK cells were incapable of lysing hiPSC-MPCs (Figures 3C-3E).

**Figure 3.**
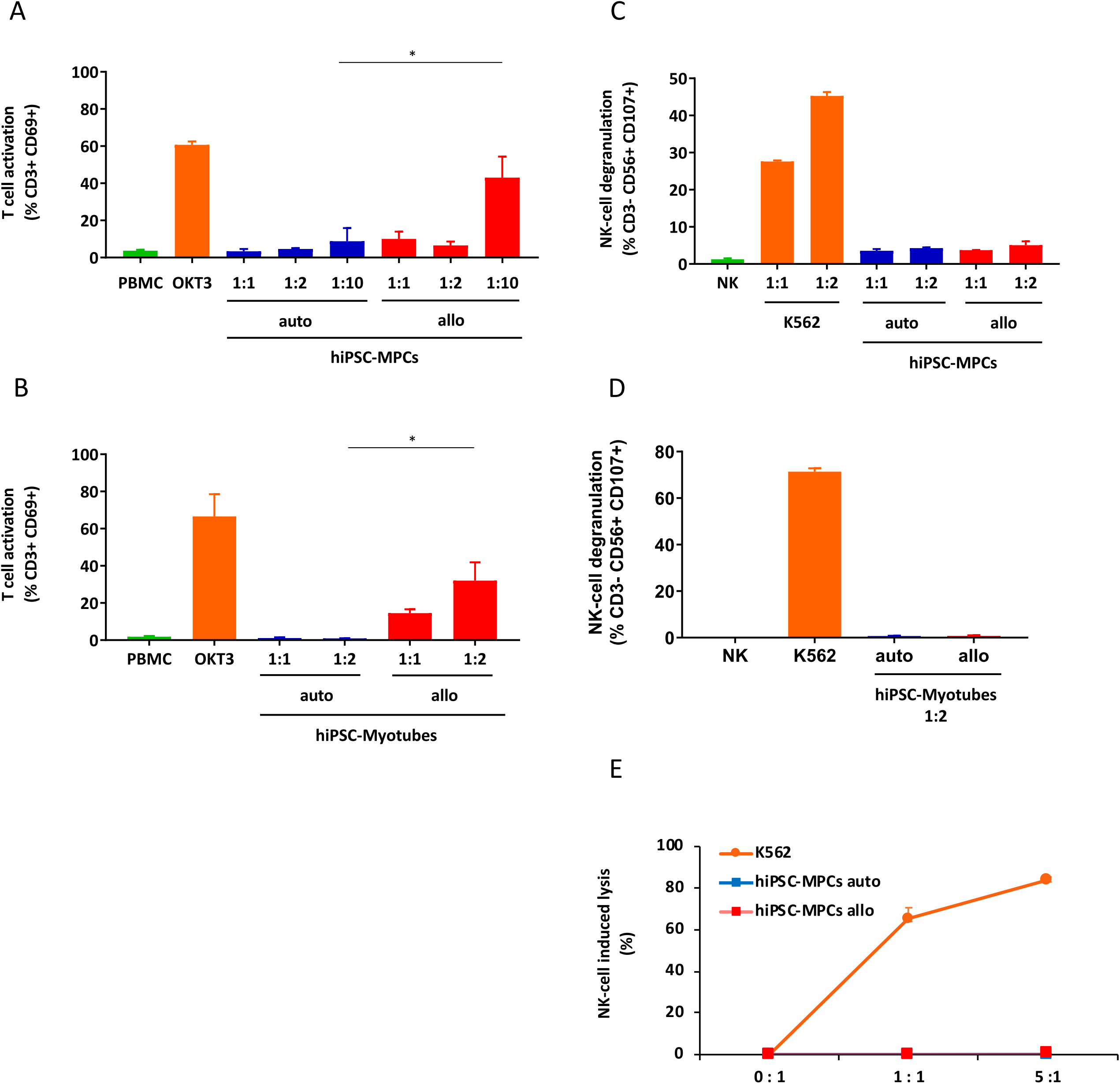
Myogenic progenitor cells derived from human iPSC are not the target of autologous immune cells *in vitro.* **(A, B)** Measures of T cell activation (as determined by gating for CD3^+^ /CD69^+^ cells) after co-culture of hiPSC-derived MPCs (panel **A**) or myotubes (panel **B**) with autologous or allogeneic PBMCs for 3 days. OKT3 was used as a positive control. Shown is the mean ± SEM of 2 independent experiments done in triplicate using cells collected from 2 different donors. **(C, D)** hiPSC-MPCs do not induce degranulation of NK cells *in vitro*. Degranulation was determined by evaluating CD107 expression on gated CD3^-^/CD56^+^ NK cell populations after a 4 hour co-culture between freshly isolated PBMCs and autologous or allogeneic hiPSC-MPCs (panel **C**) or hiPSC-myotubes (panel **D**). K562 cells were used as a positive control. Shown is the mean ± SEM of 2 independent experiments done in triplicate using cells collected from 2 different donors. **(E)** hiPSC-MPCs are not lysed by purified NK cells *in vitro*. Cell lysis was determined by flow cytometry with the absolute count of PKH26-stained hiPSC-MPCs after a 4 hour co-culture with NK cells purified from autologous or allogeneic PBMCs by magnetic negative selection. K562 cells were used as a positive control. Shown is the mean ± SEM of 2 independent experiments done in triplicate using cells collected from 2 different donors.

### hiPSC-derived myofibers are not rejected in Hu-AT mice

We next choose to further evaluate the *in vivo* immunogenicity of hiPSC-MPCs using a different humanized mouse model that combines functional T and NK cells. To this end, we adoptively transferred 1×10^7^ human PBMCs obtained from the same donors as the fibroblasts that were used to generate hiPSCs in NSG-SGM3 mice (Figure 4A). Similar to what we observed in Hu-BLT mice, more human dystrophin-positive myofibers were observed in muscles injected with autologous than allogeneic PBMCs four weeks post transplantation (Figures 4B and 4C). Muscle sections were however highly infiltrated with both allogeneic and autologous T cells (Figures 4D and 4E). Yet, the infiltration of autologous immune cells had no impact on the graft success as shown by the fact that the number of hiPSC-MPC-derived myofibers was not higher in non-reconstituted NSG mice (Figure 4C). Also, we found hiPSC-derived MPCs were not the target of NK cells *in vivo*. We believe there was a sufficient number of NK cells injected in Hu-AT mice (approximately 0.5-1,5×10^6^ calculated from the proportion of NK cells found in PBMCs) to efficiently reject MPCs based on the fact that the injection of a similar number of NK cells can prevent the growth of teratoma in the same model (Benabdallah et al. in preparation). Overall these results demonstrate that hiPSC-derived muscular grafts are immune tolerated in Hu-AT mice.

**Figure 4.**
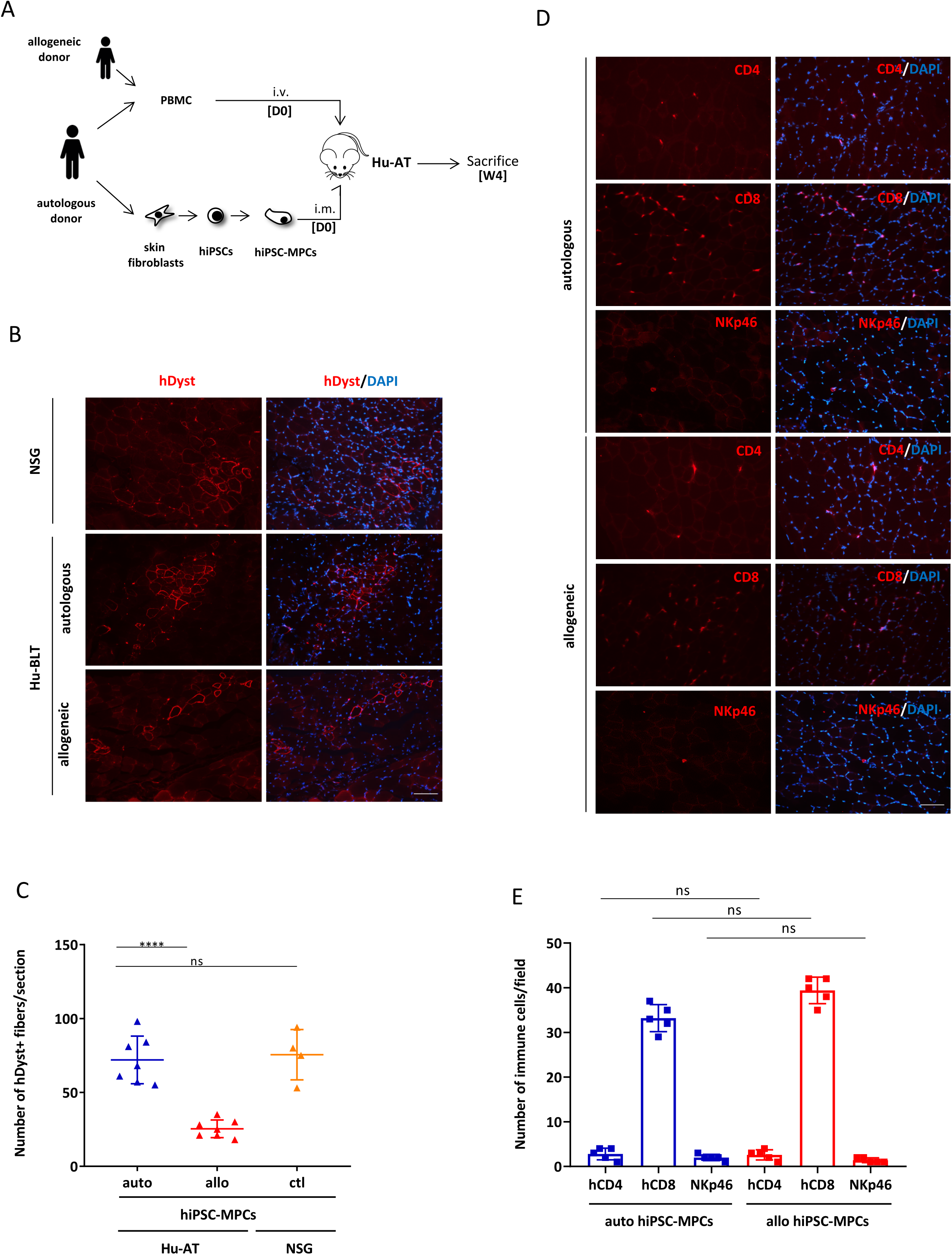
iPSC-derived myogenic progenitor cells are immune tolerated in Hu-AT mice. **(A)**Schematic of the humanized mouse model generated following the AT (Hu-AT) of human PBMCs. NSG-SGM3 mice were transplanted on day 0 (D0) with hiPSC-derived MPCs and either autologous or allogeneic PBMCs. Mice were sacrificed 4 weeks later, and skeletal muscles were harvested and muscle sections were analyzed for cell engraftment and infiltration of immune cells. **(B)** Representative photos of muscle sections from Hu-AT mice showing increased engraftment of autologous compared to allogeneic hiPSC-MPCs and resulting dystrophin positive fibers (in red). DAPI staining was performed to visualize nuclei (in blue). Showed are photos taken at 20X. Scale bar, 100 µm. **(C)** Counts of human dystrophin positive fibers observed in muscle sections of Hu-AT mice transplanted with hiPSC-MPCs as shown in panel **B**. Also showed are counts in non-immune reconstituted NSG mice to assess overall engraftment potential of transplanted cells. Each dot represents the mean ± SEM number of dystrophin positive fibers from 2-3 randomly selected sections in each muscle (n=4-7). **(D)** Representative photos showing CD4+, CD8^+^ T cell and NKp46^+^ NK cell infiltration (in red) in muscle sections of Hu-AT mice transplanted with hiPSC-MPCs and autologous or allogeneic PBMCs. DAPI staining was performed to visualize nuclei (in blue). Showed are photos taken at 20X. Scale bar, 100 µm. **(E)** Frequency of CD4+, CD8+ T cells and NKp46^+^ NK cells infiltrated in muscles of Hu-AT mice as shown in panel **D**. Each dot represents the mean ± SEM number of T cells in 2-3 randomly selected fields in each muscle (n=5).

## CONCLUSION

It will be essential to demonstrate that human iPSC-derived cells, whether obtained from autologous donors or through the development of universal cell lines ^11-13^, are tolerated by human immune cells. Using two distinct humanized mouse models, we provide evidences that MPCs are tolerated by autologous T and NK cells while they are rejected by allogeneic T cells. Moreover, the different differentiation protocols already published for the generation of MPCs from iPSCs will also need to be evaluated for their ability to evade the immune system ^14-16^. It also remains to be determined if the presence of autologous T cells would affect long-term engraftment (more than four weeks). Indeed, four weeks after the AT of PBMCs, we observed equal infiltration of T cells in both autologous and allogeneic conditions, suggesting mice were at an early stage of graft-versus-host disease. The use of immune-deficient mice that lack the murine MHC allowing to evade graft-versus-host disease but that retain T cell function upon engraftment will likely help to overcome this issue^17^. Overall, our study provides essential pre-clinical data supporting the usage of iPSC-derived MPCs in regenerative medicine.

## ACKNOWLEDGMENTS

We are grateful to the flow cytometry platform and the animal facility for providing technical support and to Renée Dicaire for handling clinical samples. This work was supported by a grant from the Canadian Institute of Health Research #MOP-126096 to C.M.B., E. H. and J.P.T., by a grant from le réseau ThéCell from the Fonds de la recherche du Québec – Santé (FRQS) and the support from la Fondation Charles Bruneau for access to technological platforms. C.M.B. was supported by a senior scientist award from the FRQS.

## AUTHOR CONTRIBUTIONS

B.B., C.D.L., M.L.G., C.C., Y.L. and C.M. performed experiments. B.B., H.E. and C.B. designed the studies, J.V.G. provided cells, J.P.T. provided cells, reagents and expertise. B.B. and C.B. wrote the manuscript.

## CONFILCT OF INTEREST STATEMENT

The authors declare no competing interests

## EXPERIMENTAL PROCEDURES

### Humanized mice

NOD/SCID/IL2Rγ null (NSG) and NSG-SGM3 (expressing human IL3, GM-CSF and SCF) mice were obtained from the Jackson Laboratory (Bar Harbor, ME) and housed in the animal care facility at the CHU Sainte-Justine Research Center under pathogen-free conditions in sterile ventilated racks. All *in vivo* manipulations were previously approved by the institutional committee for good laboratory practices for animal research (protocol #579). BLT humanized mice (Hu-BLT) were generated by surgical implantation of small pieces (1-2 mm^3^) of human fetal thymus tissues under the renal capsule and intravenous delivery of CD34^+^ hematopoietic stem cells isolated from autologous fetal liver into six week-old NSG mice previously irradiated with 2 Gy total body irradiation (1 Gy/min using a Faxitron CP-160) as previously described ^18^. Fetal tissues were obtained from consented healthy donors after surgical abortion at around week 20 of pregnancy. To monitor the human immune cell engraftment in humanized mice, peripheral blood was collected and leukocytes were purified using a red blood cell lysis solution. Cells were then labeled with conjugated antibodies for human APC-CD45, BB515-CD3, APC-H7-CD19, and PE-CD8 (all from BD Biosciences) and analyzed by flow cytometry (BD FACSCANTO II, BD Biosciences). For adoptive transfer experiments, human adult blood was collected and immune cells purified by Ficoll (GE healthcare). Mice were injected i.v. with 10 million freshly isolated PBMCs.

### Generation and characterization of hiPSCs

Fibroblasts isolated either from human fetal liver tissues or human skin after collagenase dissociation were reprogrammed into hiPSCs with the integration-free based Sendai virus (Cytotune 2.0 kit from Life Technologies). Fibroblasts were used at low population doubling to increase efficiency of reprogramming. Emerging colonies from transduced cells were manually picked and cultured under feeder-free conditions in Essential 8 medium on Geltrex-coated dishes (Life Technologies). hiPSC clones were passaged at least 15 times to increase stable pluripotency. hiPSC generation and characterization were done in the iPSC – cell reprogramming core facility of CHU Sainte-Justine. hiPSC colonies were stained with the antibodies for anti-human SSEA-4, Sox2, OCT4 and TRA1-60 overnight at 4°C using the pluripotent Stem Cell 4-Marker Immunocytochemistry Kit (Life Technologies), followed by incubation with an ALEXA secondary antibodies for 30 minutes at room temperature. Nuclei were counterstained with DAPI.

### Differentiation into myogenic progenitor cells

Myogenic progenitor cells (MPCs) were generated either directly from fibroblasts, or following fibroblast reprogramming into hiPSCs (hiPSC-MPCs). Differentiation of fibroblasts into MPCs was obtained after transduction with a MyoD-expressing adenovirus for five hours at MOI 30. hiPSC-MPCs were differentiated by first culturing hiPSC colonies in MB1 myogenic medium (Hyclone) supplemented with 10 ng/ml of bFGF for five days on Geltrex-coated culture dishes and then transduced with a MyoD-expressing adenovirus as described above ^10^. Both MPCs and hiPSC-MPCs were used the day after transduction to avoid premature fusion of the cells.

### Splenocyte and T cell activation assays

Splenocytes and PBMCs were isolated from Hu-BLT mice following mechanical digestion of the spleen or isolation using Ficoll respectively. Effector cells (splenocytes or PBMCs) were then co-cultured with either autologous or allogeneic MPCs or myotubes at 1:2 ratio during three days at 37°C. Effector cells were added to MPCs directly after transduction with MyoD or after a five day culture in 2% fetal bovine serum to allow the formation of myotubes. T cell activation was measured with a PE-conjugated anti-hCD69 on CD3^+^ gated viable cells by flow cytometry (BD LSRFortessa, BD Biosciences). Effector cells without stimulation were used as a negative control, and phytohemagglutinin (PHA) (10µg/ml) or an anti-CD3 (OKT3) antibody were used as positive controls. 7-AAD (BD Biosciences) was used to exclude dead cells.

### NK cell degranulation and cytotoxicity assays

Freshly purified NK cells (as described above) were incubated with or without target cells at the indicated ratios. K562 cells were used as a positive control in all experiments. For the NK cell degranulation assay, effector and target cells were co-cultured at 1:2 ratio in the presence of FITC-conjugated anti-human CD107a/b for one hour at 37°C, then 2µl/ml of monensin (BD Biosciences) was added to the cell mixture for an additional three hours of incubation. For cytotoxicity assay, purified NK cells (autologous or allogeneic) and PKH26-stained hiPSC-MPCs were mixed at 1:1 or 5:1 ratio and incubated for four hours at 37°C. At the end of the incubation, degranulation was quantified by flow cytometry (BD LSRFortessa, BD Biosciences) after gating on CD3-/CD56+/CD107+ viable cells and the extent of cytotoxicity was determined by the relative number of live target cells labelled with PKH26 only and dead cells labelled with both PKH26 (Sigma-Aldrich) and 7-AAD (BD Biosciences).

#### Fetal fibroblast and myoblast isolation

Fetal muscle and liver biopsies were minced into small pieces and digested using a solution of PBS with 0.2% of collagenase (Roche) and 0.25% of dispase (STEMCELL technologies) at 37°C during 30 minutes with manual intermittent mixing. Myoblasts and fibroblasts were then cultured and expanded respectively in myogenic MB1 medium and 10% fetal bovine serum-supplemented DMEM.

### Intramuscular transplantation of myogenic cells

*Tibialis anterior (TA)* muscles of mice were implanted with one million myogenic cells (either myoblasts, MPCs or hiPSC-MPCs) in 20 µl of PBS containing 10 µg/ml of cardiotoxin (Sigma). Sham mice received 20µl of PBS-cardiotoxin following the same surgical procedure. The grafted TA muscles were harvested four weeks after transplantation and frozen in optimal cutting temperature compound (OCT, VWR). Cryosections of transplanted muscles were immunostained with antibodies against human dystrophin – human specific (Developmental Studies Hybridoma Bank, University of Iowa) and human CD4 and CD8 (both from Biolegend).

#### MPC characterization and differentiation *in vitro*

MPCs cultured in MB1 were fixed and permeabilized using ethanol 95% and immunostained with a mouse anti-human desmin antibody (1:200 from DAKO) overnight at 4°C and an anti-mouse ALEXA fluor 594 (1:500 from Invitrogen) at room temperature for one hour. Confluent MPCs were maintained in Dulbecco’s modified Eagle’s medium (DMEM) supplemented with 2% fetal bovine serum and antibiotics for three-five days. Myotubes were then immunostained with a mouse anti-MyHC antibody (1:100 from the Developmental Studies Hybridoma Bank, University of Iowa) for two hours at room temperature and an anti-mouse ALEXA fluor 594 (1:500) at room temperature for one hour. Nuclei were counterstained with DAPI.

### Statistical Analysis

GraphPad Prism 8 software was used for statistical analysis; ρ values on multiple comparisons were calculated using Student’s t-Tests or One-way analysis of variance (ANOVA) with Bonferroni post Hoc test. \**p*<0.05, \*\**p*<0.01, \*\*\*\**p*<0.0001.

