## Supplementary material for "Immune tolerance of human induced pluripotent stem cell-derived myogenic progenitor cells in humanized mice": supp figures

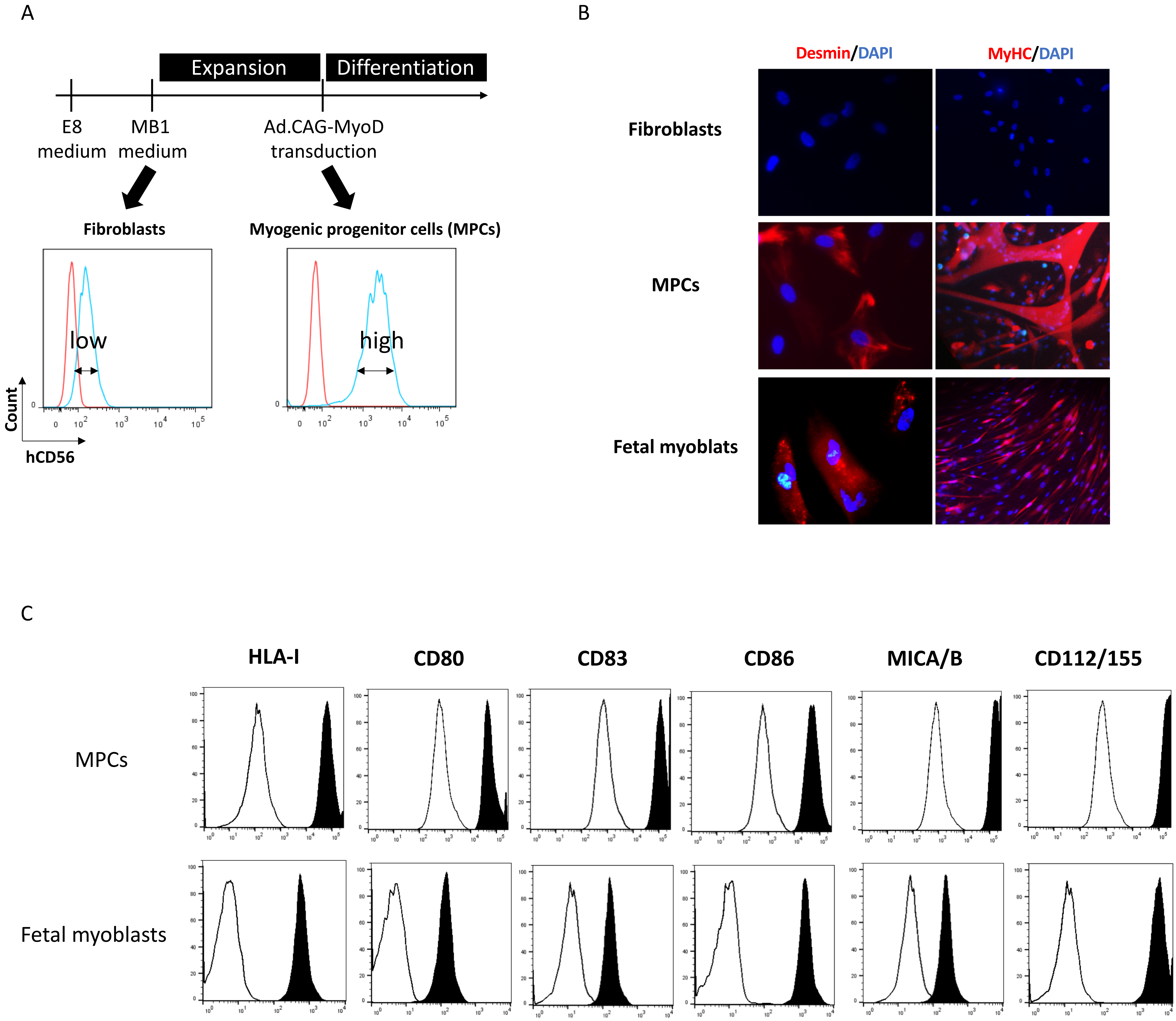


**Figure S1. Myogenic and phenotypic characterization of myogenic cells.**
(A) Schematic illustration of the protocol used for the differentiation of fibroblasts and hiPSCs in MPCs using myogenic medium (MB1) and a MyoD-expressing adenoviral vectors. Flow cytometry plots show the increased expression of the myogenic marker CD56 in differentiated cells.
(B) Representative photos showing expression of the myogenic cell marker desmin or the myosin heavy chain (in red) on MPCs and biopsy-derived fetal myoblasts compared to skin fibroblasts. DAPI staining was performed to visualize nuclei (in blue).
(C) Phenotypic characterization of hiPSC- derived MPCs and biopsy-derived fetal myoblasts. Cells were stained with the indicated mAbs (in black) or IgG isotype controls (in white) and analyzed by flow cytometry. Acquisition from one representative experiment is shown.

A

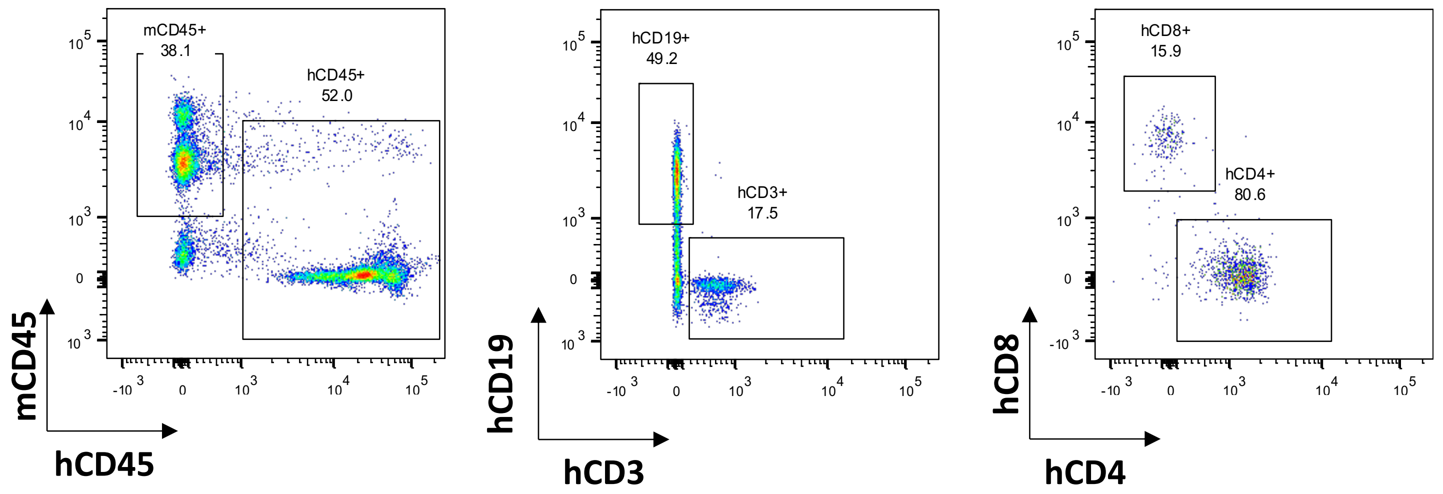


B

| Hu-BLT Mice | % hCD45 | % hCD19 | % hCD3 | % hCD4 | % hCD8 |
|---|---|---|---|---|---|
| 1 | 51 | 16 | 81 | 72 | 24 |
| 2 | 65 | 19 | 76 | 72 | 24 |
| 3 | 39 | 25 | 71 | 70 | 27 |
| 4 | 39 | 37 | 57 | 74 | 21 |
| 5 | 61 | 26 | 68 | 69 | 22 |

C

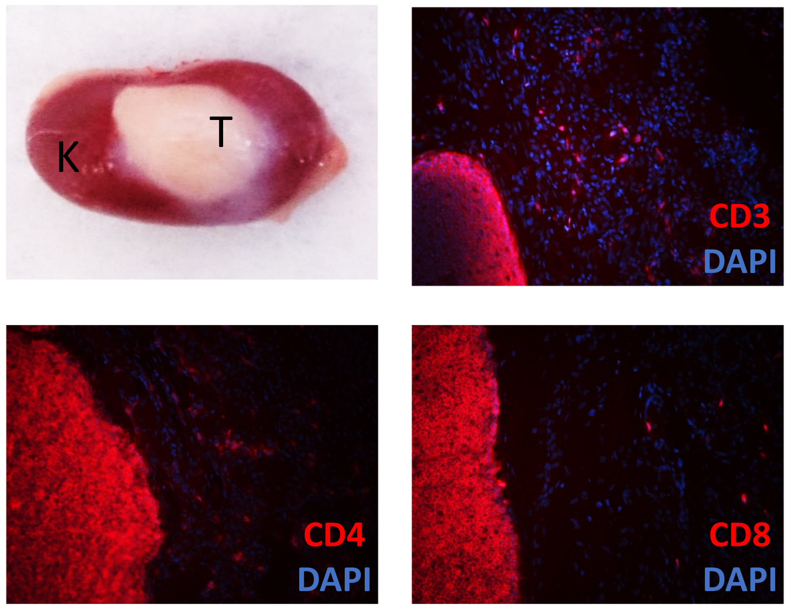


D

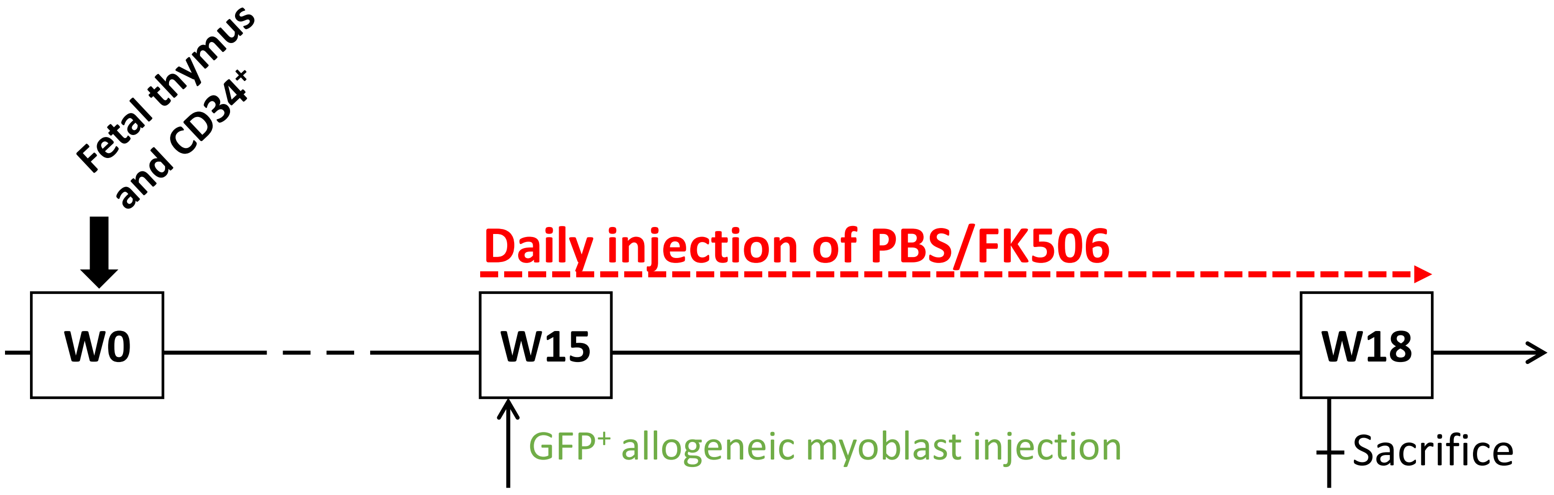


E

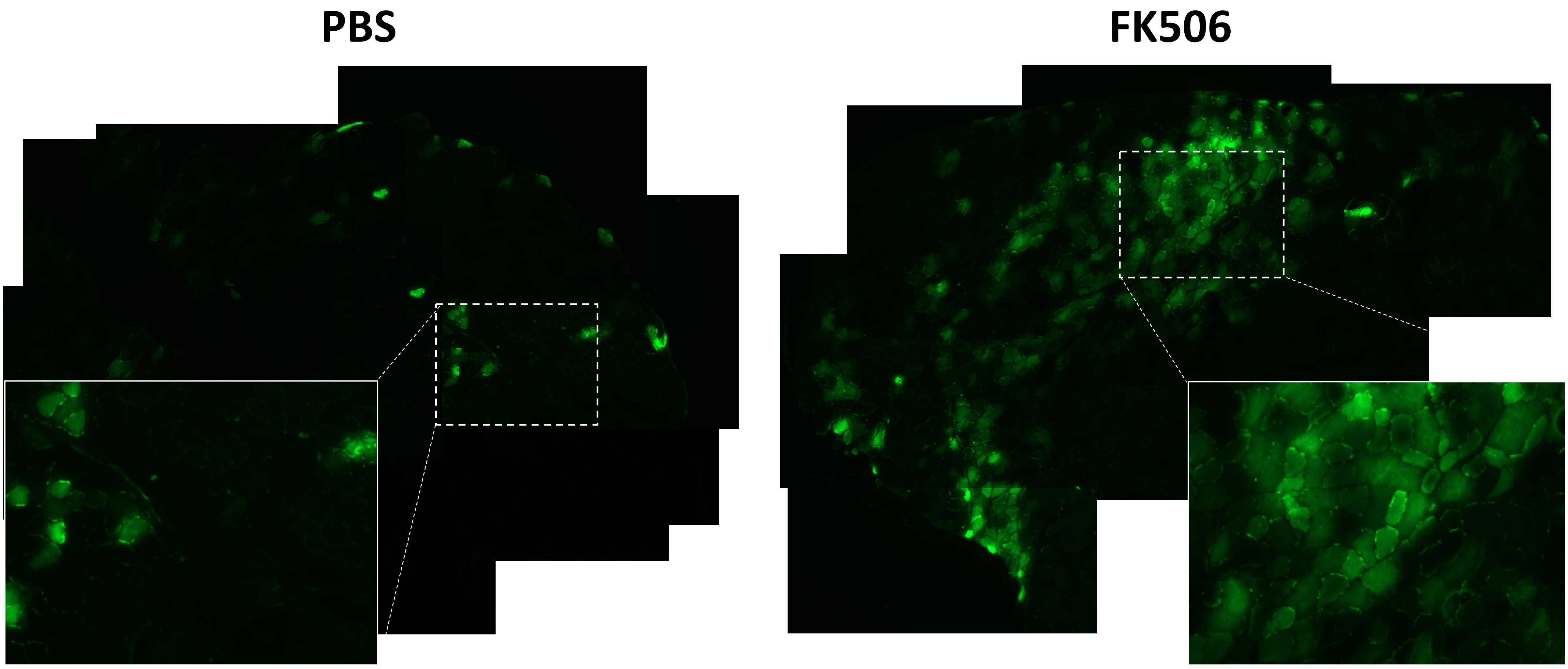

F

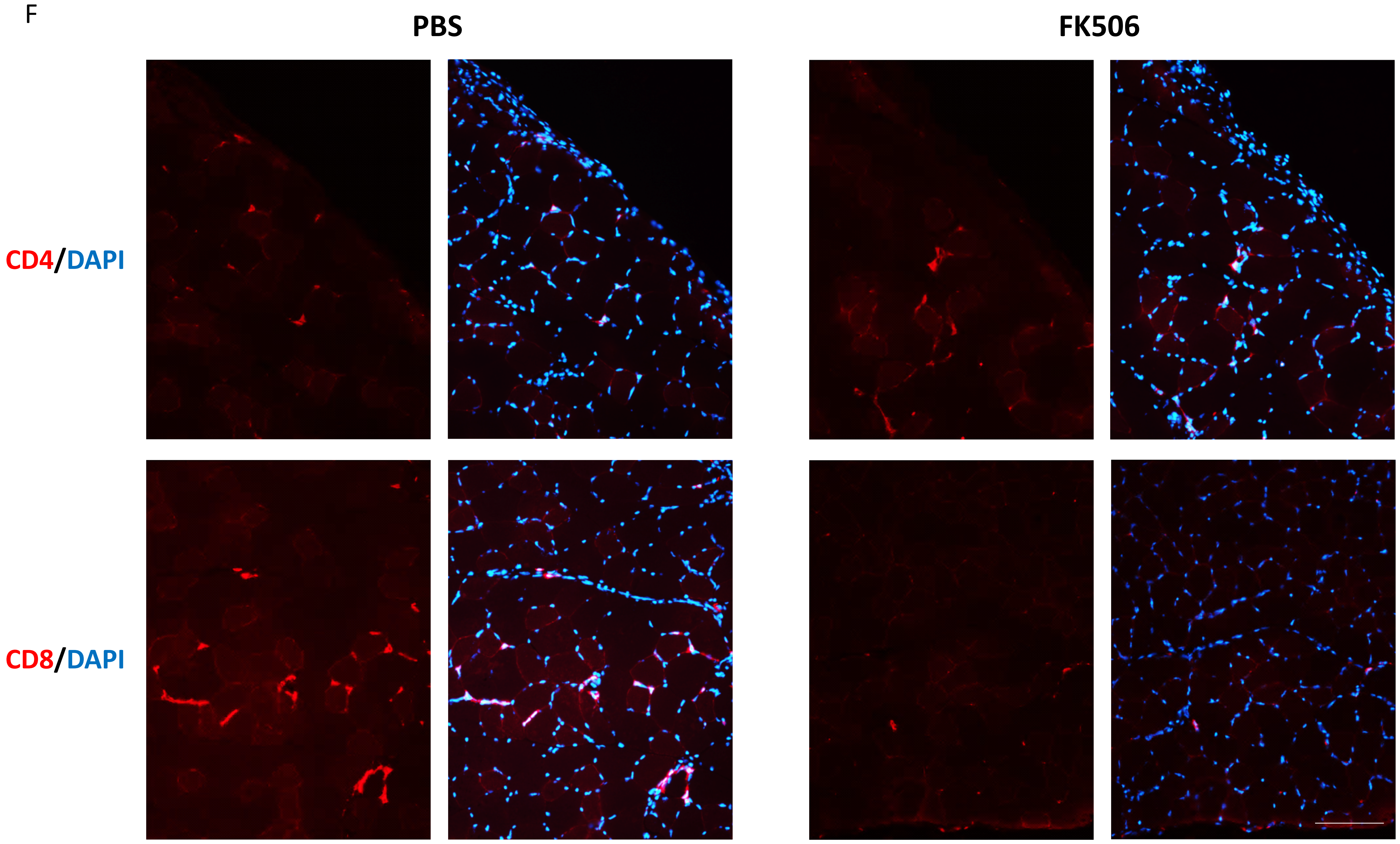


**Figure S2. Immune reconstitution in Hu-BLT mice.**
(A) Immune reconstitution of a Hu-BLT mouse 8 weeks following the transplantation of $CD34^{+}$ fetal liver cells and autologous thymic tissues. Representative plots showing human T cells (CD3, CD4 and CD8) and B cells (CD19) reconstitution in peripheral blood are shown.
(B) Frequencies of the major leucocytes subsets found in the peripheral blood of representative Hu-BLT mice 8 weeks following their reconstitution.
(C) Representative photos of a human thymic (T) implant under the mouse renal capsule (K).  Also showed are representative thymus sections showing human T cells (CD3, CD4 and CD8 in red). DAPI staining was performed to visualize nuclei (in blue).
(D) Schematic of the myoblast transplantation in the skeletal muscle of Hu-BLT mice and their immune suppression using FK506. In brief, Hu-BLT mice were generated as previously described and were injected with allogeneic myoblasts isolated from a biopsy 15 weeks (W15) post immune reconstitution. Myoblasts were modified to express the green fluorescent protein ($GFP^{+}$). Mice received daily injections of FK506 or PBS starting immediately following the injection of myoblasts until sacrifice (W18).
(E) Representative photos of the whole muscle section from Hu-BLT mice treated either with PBS or with FK506 showing increased engraftment of allogeneic myoblasts under immunosuppression as determined by the large quantity of GFP+ myofibers (in green).
(F) Representative photos showing decreased $CD8^{+}$ T cell infiltration (in red) in muscle sections of Hu-BLT treated with FK506. DAPI staining was performed to visualize nuclei (in blue). Showed are photos taken at 20X. Scale bar, 100 µm.

A

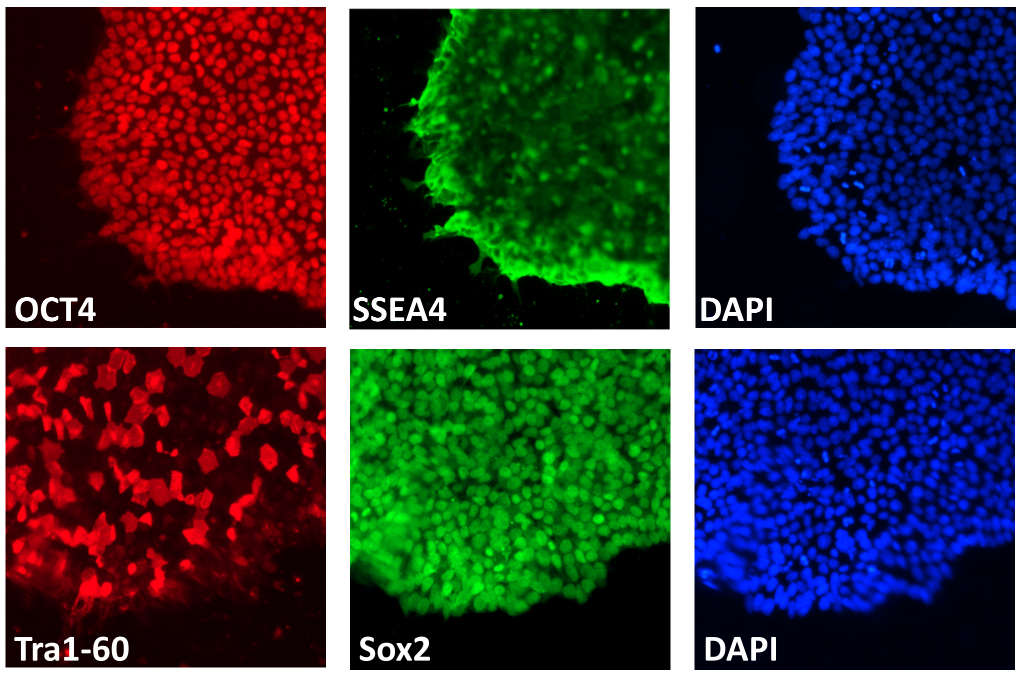


B

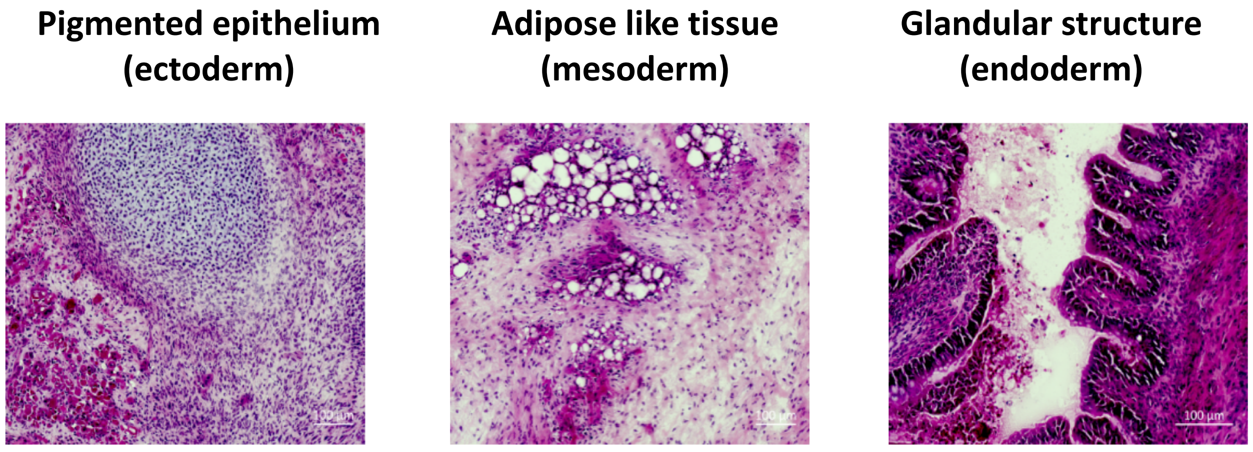


**Figure S3. hiPSC characterization**

(A) Representative photos showing the expression of pluripotency marker (Tra1-60, OCT4 in red and Sox2, SSEA4 in green) in skin fibroblast-derived hiPSCs. Cells were cultured in feeder-free conditions on Geltrex-coated dishes in E8 medium and passaged every 3-4 days.

(B) Hematoxylin and eosin staining of a teratoma-derived from 1 .$10^6$ cells of one hiPSC clone at passage 17 injected under the renal capsule of a NSG mouse and grown during 8 weeks. Representative photos showing tissues from the three embryonic germ layers are shown.
